# Large Scale Foundation Model on Single-cell Transcriptomics

**DOI:** 10.1101/2023.05.29.542705

**Authors:** Minsheng Hao, Jing Gong, Xin Zeng, Chiming Liu, Yucheng Guo, Xingyi Cheng, Taifeng Wang, Jianzhu Ma, Le Song, Xuegong Zhang

**Affiliations:** MOE Key Laboratory of Bioinformatics and Bioinformatics Division, BNRIST, Department of Automation, Tsinghua University, Beijing, China; BioMap, Beijing, China; Department of Electrical Engineering, Tsinghua University, Beijing, China; Institute for AI Industry Research, Tsinghua University, Beijing, China; Mohamed bin Zayed University of Artificial Intelligence, Abu Dhabi, UAE; School of Life Sciences and School of Medicine, Center for Synthetic and Systems Biology, Tsinghua University, Beijing, China

**Author notes:** Corresponding author: (Xuegong Zhang), (Le Song), (Jianzhu Ma). Work was done while interning at BioMap (Minsheng Hao).

## Abstract

Large-scale pretrained models have become foundation models leading to breakthroughs in natural language processing and related fields. Developing foundation models in life science for deciphering the “languages” of cells and facilitating biomedical research is promising yet challenging. We developed a large-scale pretrained model scFoundation with 100M parameters for this purpose. scFoundation was trained on over 50 million human single-cell transcriptomics data, which contain high-throughput observations on the complex molecular features in all known types of cells. scFoundation is currently the largest model in terms of the size of trainable parameters, dimensionality of genes and the number of cells used in the pre-training. Experiments showed that scFoundation can serve as a foundation model for single-cell transcriptomics and achieve state-of-the-art performances in a diverse array of downstream tasks, such as gene expression enhancement, tissue drug response prediction, single-cell drug response classification, and single-cell perturbation prediction.

## Introduction

Large-scale pretrained models are revolutionizing research in natural language processing related fields and becoming a new paradigm toward general artificial intelligence. These models trained on huge corpora become foundation models due to their fundamental importance in leading breakthroughs in many downstream tasks and their ability in discerning patterns and entity relationships within language. In life sciences, living organisms have their underlying “languages”. Cells, the basic structural and functional units of the human body, constitute “sentences” composed of a myriad of “words” such as DNA, RNA, proteins, and gene expression values. An intriguing question is: Can we develop foundation models of cells based on massive cell “sentences”?

Single-cell RNA sequencing (scRNA-seq) data, also known as single-cell transcriptomics, offers high-throughput observations into cellular systems^1–3^, providing massive archives of transcriptomic sentences of all types of cells for developing foundation models of cells. In transcriptomic data, gene expression profiles depict complex systems of gene-gene co-expression and interaction within cells. With the efforts of the Human Cell Atlas^4^ and many other studies, the data scale is exponentially growing^5^. With about 20,000 protein-coding genes across millions of cells, the observed gene expression values scale to a magnitude of trillion “tokens” (Table S1), which is comparable to the volume of natural language texts used to train large language models (LLMs) such as GPT. This provides the foundation for us to pre-train a large-scale model to extract complex, multifaceted internal patterns of cells in a manner similar to LLMs learning human knowledge from massive archives of natural language texts.

In the LLMs pre-training^6,7^, the growth in both model and data scale is critical for constructing a foundation model that can effectively mine intricate multi-level internal relationships. Recent efforts have been reported on pre-training models on single-cell gene expression data^8–11^, but the development of large-scale foundation models still presents unique challenges. First, the gene expression pre-training data needs to encompass a landscape of cells across different statuses and types. Currently, most scRNA-seq data are loosely organized, and a comprehensive and complete database is still lacking. Second, when modeling each cell as a sentence and each gene expression value as a word, the nearly 20,000 protein-coding genes make the “sentence” exceptionally long, a scenario that traditional transformers struggle to handle^12,13^. Existing work had to restrict their models to a small list of selected genes. Third, scRNA-seq data across different sequencing techniques and laboratories exhibit high variance in sequencing read depth. This technical noise prohibits the model from learning uniform and meaningful representations for cells.

In this study, we address these challenges and designed the first large-scale foundational model scFoundation of 100M parameters working on ∼20,000 genes. We collect the largest scRNA-seq data set with over 50 million gene expression profiles for pre-training, covering cells from different statuses and various tissues. We develop an asymmetric architecture designed for single-cell RNA-seq data to accelerate the training process and improve model scalability. We develop a read-depth-aware pre-training task that enables scFoundation not only to model the gene co-expression patterns within a cell but also to link the cells with different read depths.

To verify the ability of scFoundation for learning the characteristics of both cells and genes, we conduct experiments on multiple downstream tasks, including gene expression enhancement, drug response prediction on bulk data, single-cell drug response classification, and single-cell perturbation prediction. scFoundation achieved state-of-the-art performance by adapting context embeddings to the corresponding downstream models. Our work reveals the efficacy and value of large-scale pre-trained models for transcriptomics data and demonstrates its foundation function in facilitating both biology and medical task learning. We explore and push the boundaries of the foundation model in the single-cell field.

## Results

### The scFoundation pre-training framework

We developed a large-scale model, scFoundation, modeling 19,264 genes with 100 million parameters pre-trained on over 50 million scRNA-seq data. To the best of our knowledge, this is the largest model parameter size, gene coverage, and data scale in the single-cell field. The ability to efficiently train such a model benefited from our pre-training framework, which consists of three parts: model design, pre-training tasks, and data collection (Fig. 1A).

**Figure 1.**
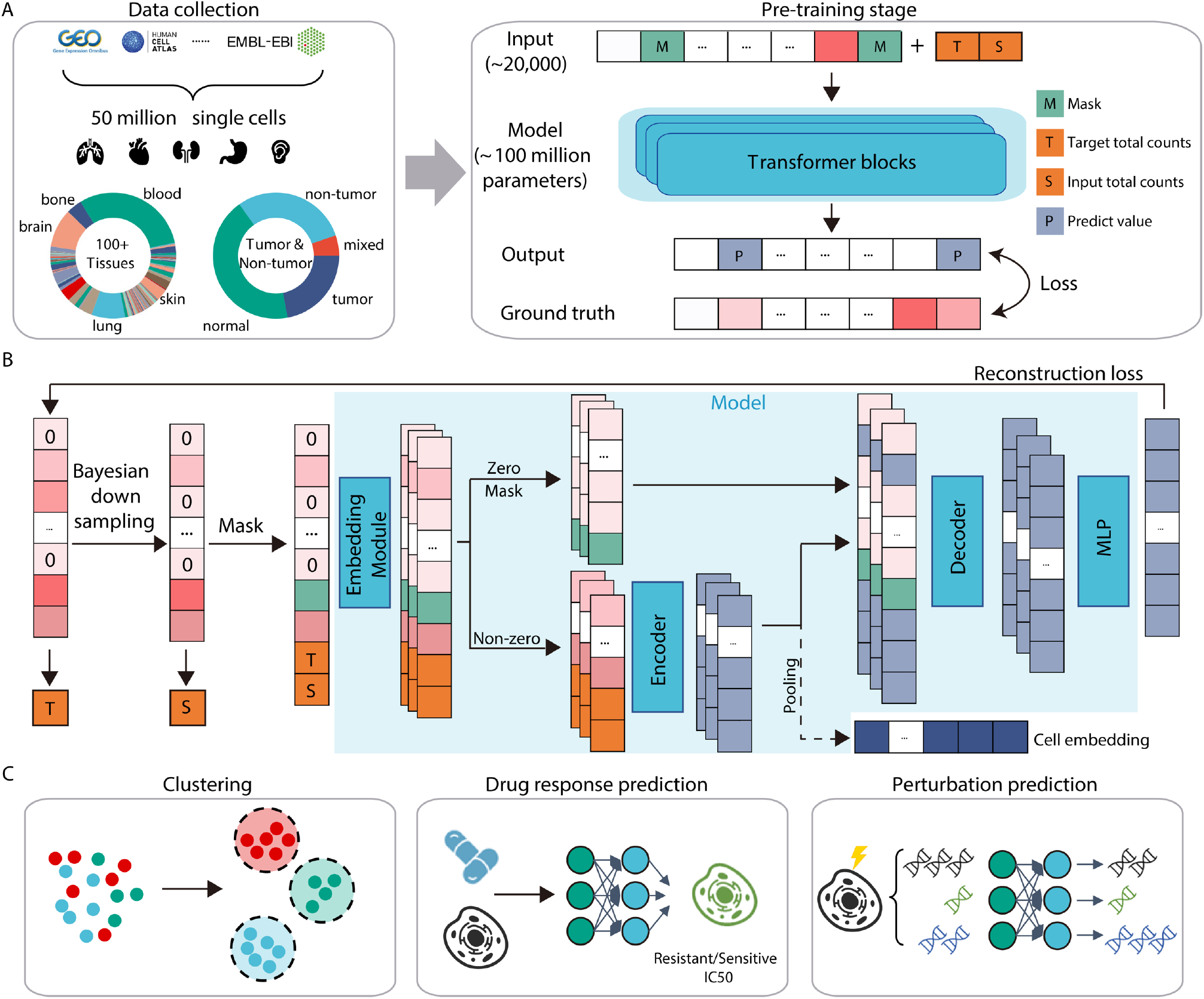
The schematic overview of the pre-training framework. A) 50 million single-cell gene expression profiles were collected, covering tumor and non-tumor cells from various tissues. These data were used for the read-depth-aware (RDA) task to pre-train the model. In the RDA task, the input consists of the masked gene expression vector and two total count indicators (T and S). The output is the predicted expression value for all genes, and the loss is computed at the masked positions. B) Outline of the pre-training process. A raw gene expression vector serves as a training sample. A hierarchical Bayesian downsampling strategy generates the input sample. The gene expression total count values (T and S) of the raw and input samples are computed. Values in the input sample are randomly masked. The scalar values are converted into embeddings. Only embeddings corresponding to non-zero and non-masked values (including T and S) are fed into the model encoder. The output embeddings of the encoder are then combined with mask and zero embeddings and fed into the decoder. Also, the encoder output can be pooled to generate a cell embedding for downstream usage. The decoder output embeddings are projected to the gene expression value via a shared multilayer perceptron (MLP) layer. The regression loss between the predicted and raw sample’s gene expression values is computed. C) The pre-training embeddings can be leveraged as substitutes for the gene expression profiles, facilitating read depth enhanced cell clustering, drug response prediction, and perturbation prediction.

We developed xTrimoGene, a scalable transformer-based model with both algorithmically efficient and engineering acceleration strategies^14^. It included an embedding module and an asymmetric encoder-decoder structure. The embedding module converted continuous gene expression scalars into learnable high-dimensional vectors, which were then used as the input of the encoder and decoder. This module fully retained information from the raw expression values, a notable improvement over the discretized values used in previous models^9,15^. The asymmetric encoder-decoder architecture was specifically designed to accommodate the high sparsity characteristics of single-cell gene expression data. The encoder only accepted embeddings of the non-zero and non-masked expressed genes as input, and the decoder accepted all genes’ embedding (i.e. the encoder processed embeddings and zero and mask embeddings) as input. This architecture gave differential attention and computational resources to zero and non-zero values, thereby achieving the efficient learning of all gene relationships without any selection (e.g., highly variable gene selection). Moreover, the model deployment incorporated a variety of large-scale model training optimization techniques to ensure efficient training (Methods).

We developed a new pre-training task called the read-depth-aware (RDA) modeling, which was an extension of masked language modeling^16^, considering single-cell gene expression data had a high variance in read depth, especially in the large-scale pre-training data. In RDA, we trained the model to predict the masked gene expression of a cell based on other genes’ context. The context was from a counterpart or a low read-depth variant of that cell’s gene expression profile. Specifically, we processed a raw gene expression training sample with a hierarchical Bayesian downsampling strategy to generate an input sample with an unchanged or altered total count (Methods). We treated the total count of all gene expressions as the measure of one cell’s read depth and defined two total count indicators: T (representing ‘target’) and S (representing ‘source’), corresponding to the total counts of the raw and input samples respectively. We randomly masked the expression values of genes in the input sample and recorded the index of masked genes. Then the model took the masked input sample and two indicators to predict the expression value of the raw sample at the masked index (Fig. 1B). A regression loss was conducted on the predicted and raw gene expression values. This pre-training process enabled the pre-trained model not only to capture the gene-gene relationship within the cell but also to harmonize the cell with different read depths. When used for inference, the number T could control the total count of the output gene expression profiles. Thus, we could feed the cell’s raw gene expression to the pre-training model and set the T higher than its total count S to generate a gene expression value with enhanced sequencing depth.

We constructed a comprehensive single-cell gene expression data set by collecting data from all publicly available single-cell resources, including GEO^17^, Single Cell Portal, HCA^4^, EMBL-EBI^18^, etc. We aligned all data to a consolidated gene list that comprised 19,264 protein-coding and common mitochondrial genes, as identified by the HGNC^19^. After data quality control (Methods), we got over 50 million human scRNA-seq data for pre-training. The abundant data sources made our pre-training dataset rich in biological patterns. From an anatomical perspective, the data covered more than 100 tissue types under different diseases, tumors, and normal states (Fig. 1A); From a cell ontology perspective, the data encompassed almost all known human cell types and cell states.

After pre-training, we applied the scFoundation model to multiple downstream tasks (Fig. 1C). The outputs of the scFoundation encoder were pooled into cell embeddings, which were used for cell clustering and drug response prediction. The outputs of the scFoundation decoder were gene-level context embeddings, which were used for perturbation prediction.

### scFoundation is a scalable model for read-depth enhancement without fine-tuning

In LLMs, researchers observed a power-law relationship between the loss and various factors, including model size, the amount of computation utilized during training, etc^6,20^. This relationship, commonly referred to as the scaling law, was also confirmed in our framework. Fig. 2A illustrated the training of three models with parameter sizes of 3 million, 10 million, and 100 million. It was found that as the model parameters and the total number of floating-point operations (FLOPs) increased, the loss on the validation dataset exhibited a power-law decline. We then estimated the performance of various scale xTrimoGene architecture models with parameter size equivalent to previous models^8–10,15^. Our scFoundation model, with 100 million parameters, surpassed all these models in terms of parameter size and achieved the lowest reconstruction loss.

**Figure 2.**
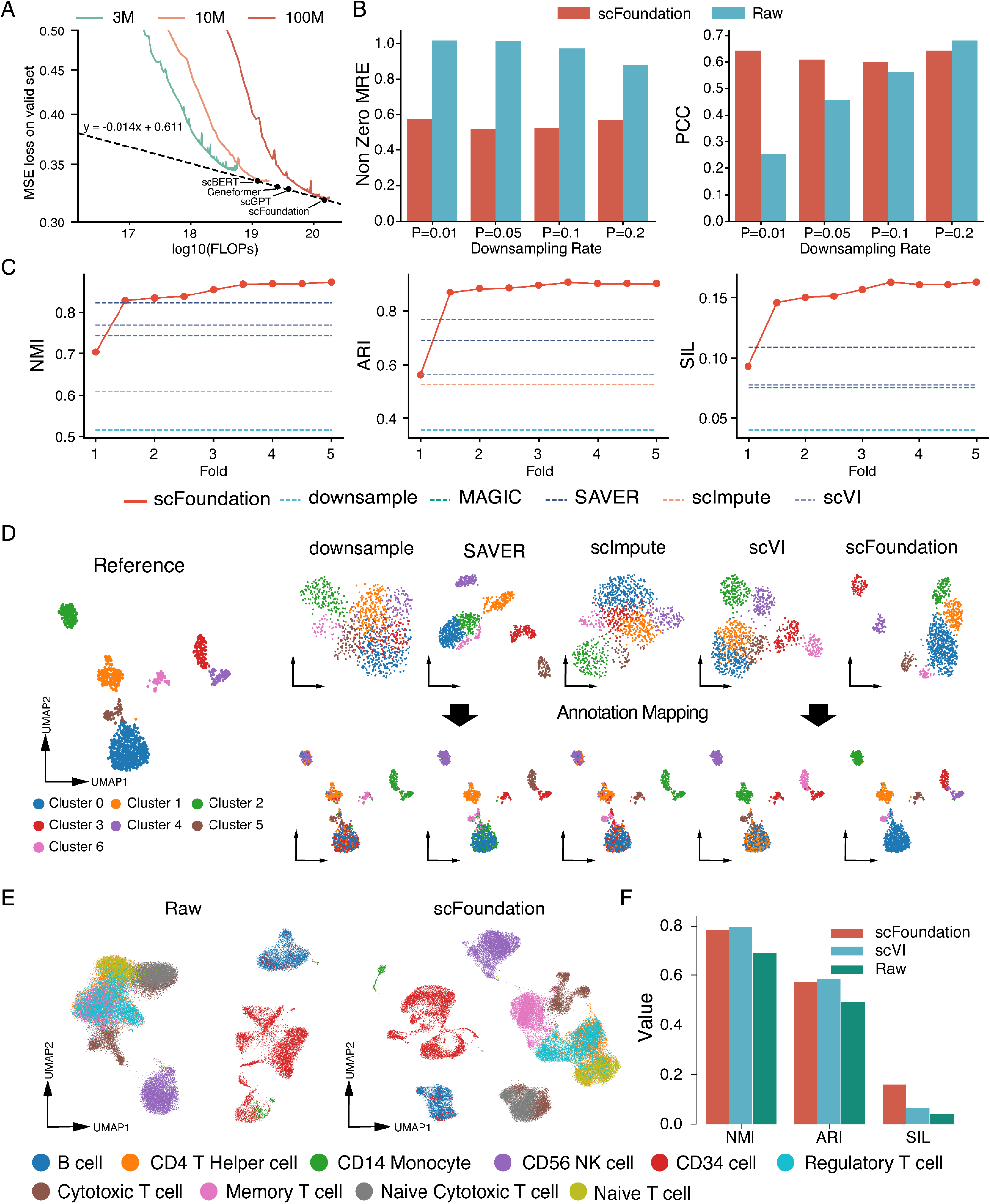
A) Training loss under different parameter sizes and FLOPs. The dots noted as previous models are xTrimoGene architecture models with parameter size equivalent to their scale. B) Evaluation of read depth enhancement performance on an unseen dataset. Mean relative error among non-zero expressed genes and Pearson correlation among all genes were used as metrics to evaluate the recovered gene expression performance. Lower error values indicate better performance, while higher Pearson correlation values indicate better performance. C) Comparison of the scFoundation model with other imputation methods based on cell clustering metrics. The x-axis represents the fold change between the desired total count value and the input total count value, while the y-axis represents the score. D) UMAP plots of cell embeddings generated by different methods. The left plot shows the reference UMAP plot obtained using raw gene expression, with colors indicating cell clusters. The upper-right plots display clustering results obtained by different methods (Sample, MAGIC, scImpute, scVI, and scFoundation). “Sample” refers to no imputation process. The number of clusters is aligned. The lower-right plots depict the clustering results of each method mapped onto the reference UMAP plot. E) UMAP plot comparing raw gene expression and scFoundation-imputed cell embeddings on the Zheng68K dataset. F) Comparison of clustering performance among scFoundation, scVI, and raw data on the Zheng68K dataset.

Our RDA modeling enables scFoundation to enhance the read depth of the input cell by setting T as a higher number than S. We first assessed this ability on an independent test dataset. We downsampled the total count into 1%, 5%, 10%, and 20% of the original profiles, generating four corresponding datasets with varying total count fold changes. For each dataset, we didn’t finetune the scFoundation but directly utilized it to enhance the cells with a low total count by setting the desired total count T as the reciprocal of the sample rate. We computed the mean absolute error (MAE) and mean relative error (MRE) metric between the predicted and ground-truth values of non-zero expressed genes (refer to Methods). Additionally, we evaluated the Pearson Correlation Coefficient (PCC) of gene expression between the enhanced and ground-truth data. We employed the downsampled gene expression values as the baseline for comparison. As depicted in Fig. 2B and Fig. S1, scFoundation demonstrated a significant reduction of half the MAE and MRE compared to the non-enhanced baseline. Particularly, when the downsampling rate was below 10%, the baseline results exhibited a 97.5% MRE, indicating that most of the non-zero values were sampled as zeros. However, scFoundation consistently maintained an MRE of 60%. As the downsampling rates increased, the baseline results showed a lower error and a higher PCC, yet scFoundation continued to outperform the baseline. These findings highlight the ability of scFoundation to enhance gene expressions in scenarios even with extremely low total counts, and robust to be generalized to new datasets.

Clustering is another common analysis to validate the performance of read-depth enhancement methods. We compared scFoundation with imputation methods including MAGIC^21^, SAVER^22^, scImpute^23^, and scVI^24^ on a human pancreatic islet dataset^25^ processed by SAVER. This dataset comprised manually generated downsampled gene expression profiles and their corresponding reference data. For scFoundation, we set T as the different folds of S (ranging from 1 to 5), fed the downsampled data, and obtained the five sets of cell embeddings. For other methods, we first used the downsampled data to train the methods. Then we got cell embeddings from scVI and got the imputed gene expression from other methods. The ground truth cluster labels were obtained from the reference data. For evaluating clustering accuracy, we employed metrics including Normalized Mutual Information (NMI), Adjusted Rand Index (ARI), and Silhouette Coefficient (SIL). The clustering performance obtained from the downsampled data was used as the baseline.

We found that scFoundation already outperformed both the baseline and scImpute in all metrics when T was equal to S (fold change=1, Fig. 2C), indicating that the scFoundation embeddings provided more informative representations compared to the raw gene expression. As the fold change increased, we observed an initial “jump” in scFoundation’s performance, surpassing the other methods, followed by a continuous improvement in all metrics. We generated Uniform Manifold Approximation and Projection (UMAP) plots for visualizing the scFoundation embedding results at a fold change of 5 and the results obtained from other methods (Fig. 2D). Notably, scFoundation’s cell embeddings exhibited distinct cluster boundaries compared to the baselines and other methods. Furthermore, we performed clustering on all methods’ results and mapped the cluster labels onto the reference UMAP. The labels produced by the other methods were mixed in the reference, particularly within the ground truth cluster 0. scFoundation stood out as the only method that visually demonstrated aligned cell cluster assignments consistent with the reference results.

We then applied scFoundation to the Zheng68K dataset from human peripheral blood mononuclear cells (PBMC)^26^. This dataset had about 600,000 cells, comprised of cell types that exhibited high similarity and were sequenced using the first version of the 10X Chromium platform. Each cell in this dataset only had non-zero expression values for approximately 500 genes, and the total count was less than 2000 (Fig. S2). We fed these cells into scFoundation to obtain the read depth enhanced cell embeddings by setting T as 10,000.

Comparing the UMAP plot of the scFoundation embeddings with that of the raw gene expression profiles, we observed that the scFoundation embeddings made it easier to distinguish Memory T cells from other T cell populations and exhibited improved discrimination between CD14 Monocytes and CD34 cells. To further evaluate the clustering performance with other methods, we trained the only scalable model scVI on this dataset and compared it with the scFoundation embeddings (Fig. 2F). Both clustering results outperformed the raw gene expression profiles and achieved comparable performance in terms of NMI and ARI metrics, and the scFoundation embeddings had a higher SIL score.

These results demonstrated that scFoundation possessed the capability to enhance the read depth of cells. Notably, an important distinction between scFoundation and other imputation methods was that scFoundation could achieve the best performance without the need for dataset-dependent training procedures.

### scFoundation improves cancer drug response prediction

Cancer Drug Responses (CDRs) study the tumor cell response upon drug intervention. Computationally predicting CDR is critical to guiding anticancer drug design and understanding cancer biology^27^. We combined scFoundation with a CDR prediction method, DeepCDR^28^, to predict the half-maximal inhibitory concentrations IC50 values of drugs across several cell line data. This experiment served as a validation of whether scFoundation could provide informative embeddings for bulk-level gene expression data, despite being trained on single cells.

The original DeepCDR model used drug structural information and multi-omics data as input and outputted the predicted IC50. Here, we focused on gene expression data and replaced the transcriptome MLP subnetwork in DeepCDR with scFoundation. In other words, we used scFoundation to extract transcriptome features and fed them into the subsequent prediction module (Fig. 3A). We integrated the Cancer Cell Line Encyclopedia (CCLE)^29^ and Genomics of Cancer Drug Sensitivity (GDSC)^30^ datasets to obtain the input cell line gene expression data, the input drugs and IC50 labels.

**Figure 3.**
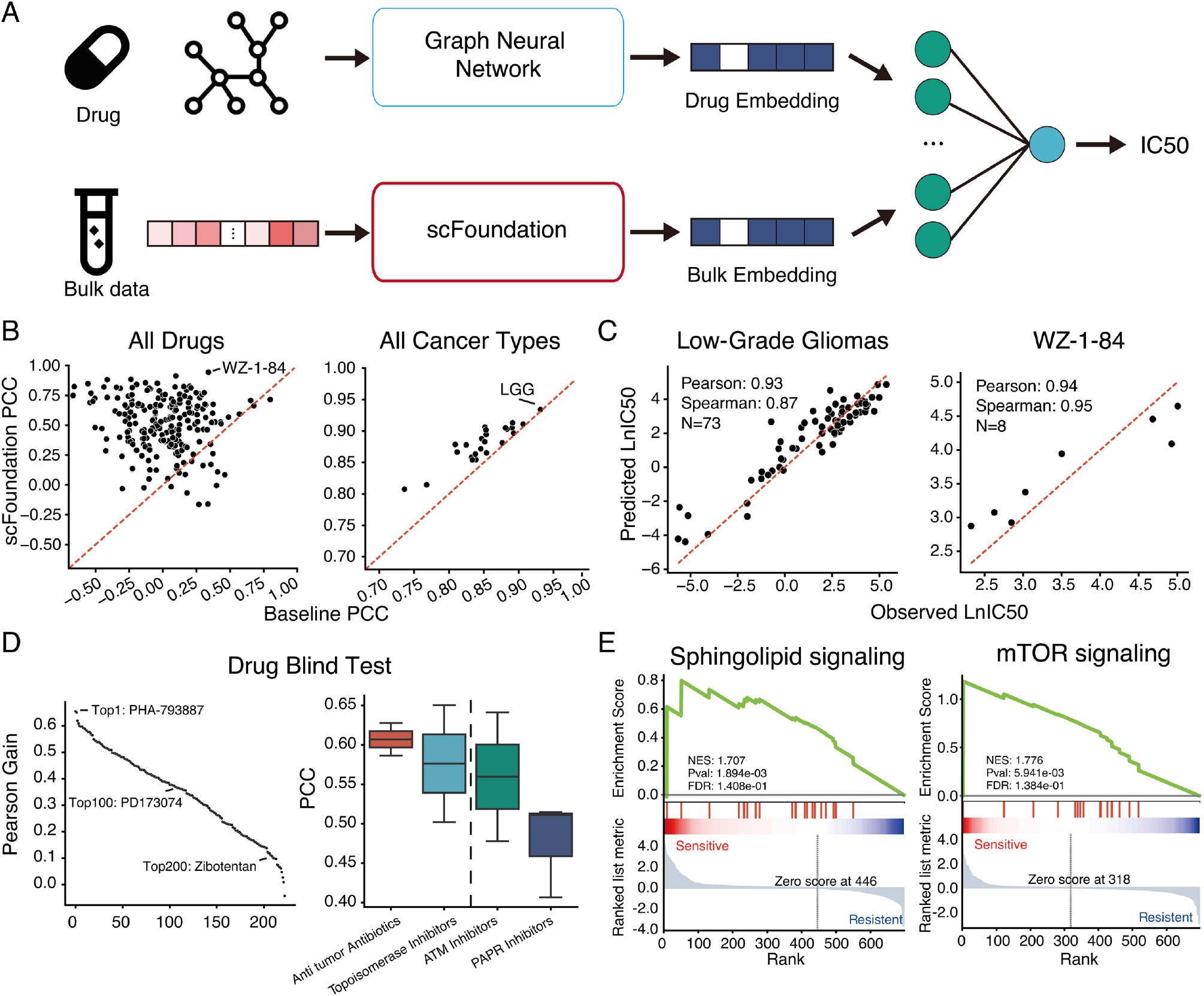
Performance of drug response prediction using scFoundation embeddings. A) Illustration of the embedding-based CDR prediction model. B) Pearson correlation coefficient (PCC) between all drugs and cancer types in the test set. Each dot represents a drug or cancer type, with the x-axis and y-axis showing the PCC obtained by DeepCDR using gene expression and scFoundation embeddings, respectively. C) Comparison of predicted and observed IC50 values for the drug WZ-1-84 on the cancer-type low-grade gliomas. Each dot represents a drug and cell line combination. D) Leave-one-drug-out blind test performance. The left plot shows the PCC gain obtained by replacing gene expression with cellular embeddings. Each dot represents a drug, with the y-axis indicating the gained PCC values and the x-axis representing the rank. Higher-ranked drugs have a higher PCC gain. The right plot groups the drugs into different types. E) Gene set enrichment analysis (GSEA) results on cell line data with lower predicted IC50 values. The Sphingolipid signaling pathway was enriched in Doxorubicin-sensitive cell lines, while the mTOR signaling pathway was enriched in Vorinostat-sensitive cell lines.

We evaluated the performance of scFoundation-based results and gene expression-based results across multiple drugs and cell lines (Fig. 3B). Most drugs and all cancer types achieved a higher PCC by using scFoundation embeddings. We further visualized the best prediction case of drug and cancer types (Fig. 3C). Regardless of the high or low lC50, the scFoundation embedding-based DeepCDR model could predict accurate values and achieved a PCC above 0.93.

The CDR prediction models were often used to predict the IC50 of the new unseen compound across the cell line. For testing the generalization ability to new drugs, we then left one drug-related dataset at a time as the test set and used the remaining data as the training set. Under this drug-blind test, we found that models based on scFoundation embeddings consistently outperformed the original model. The top 1 PCC-gaining drug, PHA-793887, was a potent ATP-competitive CDK inhibitor, and its PCC improved from 0.07 to 0.73. Even for the 200th-ranked drug, Zobotentan, which was used for blocking Endothelin A receptor activity, its PCC improved from 0.49 to 0.64.

We further grouped drugs into different therapy types to examine whether the IC50 prediction performance was related to their intrinsic mechanisms. We observed that, based on scFoundation-predicted results, drugs belonging to chemotherapy, such as anti-tumor antibiotics and topoisomerase inhibitors, had a higher PCC than drugs belonging to targeted therapy, such as ATM and PARP inhibitors (Fig. 3C right). This may be due to the fact that specific gene mutations often have an important impact on targeted therapy^27^, but mutation information is difficult to reveal from gene expression data; while chemotherapy drugs were widely reported to be related to gene expression^31,32^ so their IC50 is easier to predict. Gene expression-based results had an overall lower PCC, and we did not observe a significant difference between drug types.

Then we used our model to predict unknown CDR in the data, and thus each drug would have a predicted IC50 on all cell lines. To validate these new predictions, we performed a Gene Set Enrichment Analysis (GSEA)^33^ on the new predictions with relatively low IC50, which indicated the sensitivity of the cell line to the drug. For instance, the sphingolipid signaling pathway was enriched in Doxorubicin-sensitive cell lines. According to the KEGG database^34^, this pathway was related to sphingomyelin (SM) and its metabolism. And SM was reported to interact synergistically with Doxorubicin by altering cell membrane permeability resulting in a lower IC50 of the drug in these cell lines^35^. Additionally, the mTOR signaling pathway was enriched in Vorinostat-sensitive cell lines. Previous studies have shown that Vorinostat inhibits carcinoma growth by dampening the mTOR signaling pathway^36^. And other clinical studies have also shown that mTOR inhibitors were often used in conjunction with Vorinostat^37,38^, suggesting the relationship between Vorinostat and the mTOR pathway. These examples support the validity of our predictions.

Our results demonstrated that although the scFoundation model was pre-trained on single-cell transcriptomics data, the learned gene relationships were transferable to bulk-level expression data to produce condensed embeddings, facilitating more accurate IC50 prediction. These findings illustrated the potential of scFoundation in expanding our understanding of drug responses in cancer biology and possibly guiding the design of more effective anticancer treatments.

### scFoundation transfers bulk-level drug sensitivity prediction model to single-cell data

Intratumor heterogeneity is a well-known issue that often leads to treatment failure^39,40^. Inference of drug sensitivities at the single-cell level can help identify specific cell subtypes that exhibit different drug resistance characteristics, offering valuable insights into underlying mechanisms and potential new therapies^41–43^. We applied scFoundation to tackle this crucial task of single-cell level drug response classification based on the SCAD^44^ downstream model. Due to the fact that the limited availability of cellular-level drug response data for only a small subset of drugs and cancer types, SCAD was trained on the bulk-level datasets to learn pharmacogenomic information and then transferred this knowledge to infer drug efficacy at the single-cell level. In other words, it took both bulk and single-cell data, affected by the same drug, as input and provided sensitive or non-sensitive labels for each cell. In our study, we used scFoundation to obtain unified embeddings of bulk and single-cell data (Fig. 4A). Subsequently, we used these embeddings to train SCAD models and assessed the classification performance.

**Figure 4.**
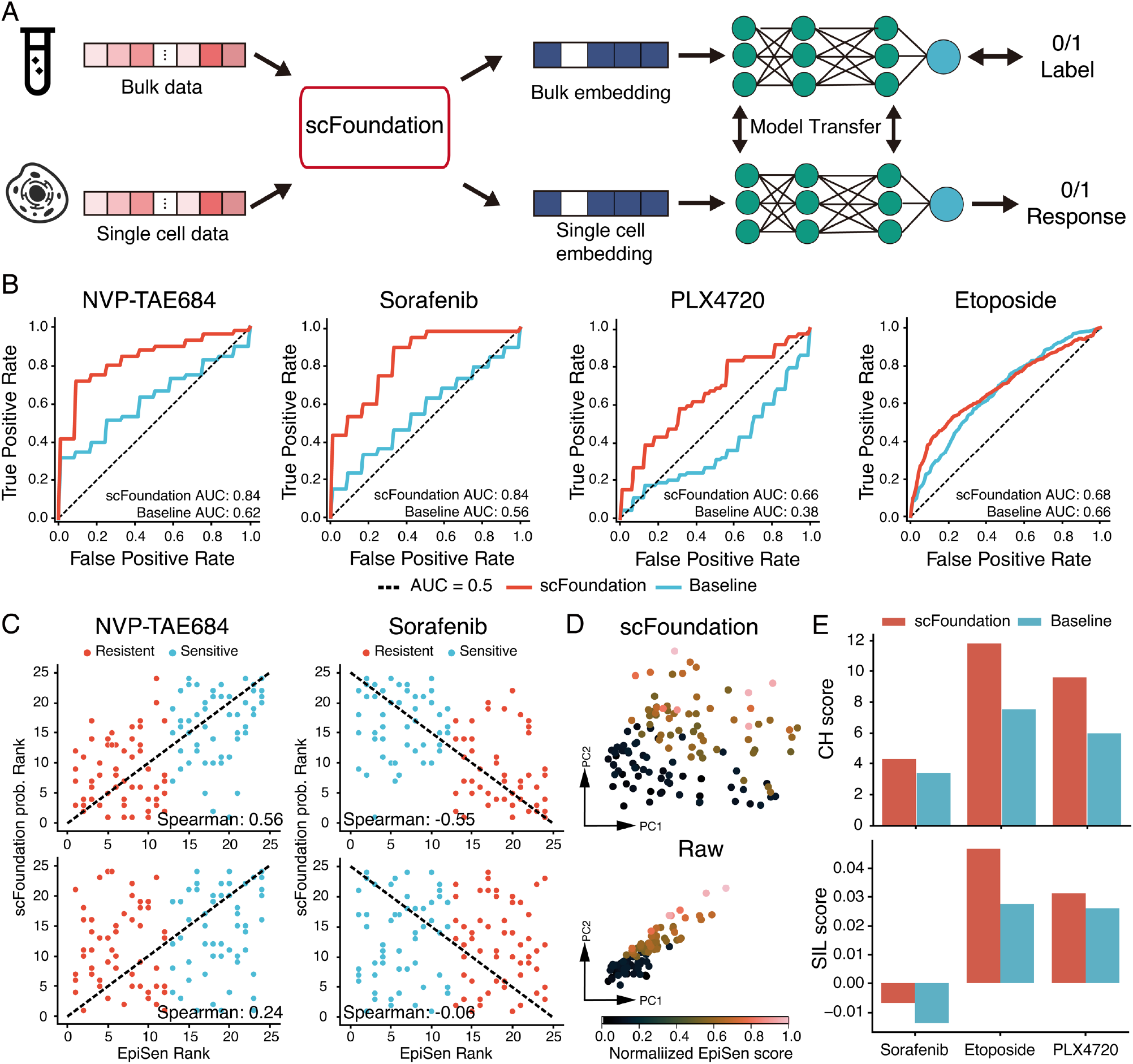
Single-cell drug response classification tasks based on scFoundation cell embeddings. A) Illustration of the embedding-based single-cell response classification model. B) Receiver operating characteristic (ROC) curves for the four drugs. The red and blue lines represent the performance of SCAD using scFoundation embeddings and gene expressions, respectively. C) Correlation between drug sensitivity probability and normalized EpiSen score. Each row corresponds to a model, and each column represents a drug. D) PCA plot of the SSC47 single-cell dataset from cell lines. Color denotes the reference epithelial senescence-related program (EpiSen) score. Cells with different EpiSen scores exhibit distinct responses to drugs. E) Clustering performance on all three drugs-related bulk datasets. Each bulk dataset has two types of labels: sensitive and resistant. CH score: Calinski-Harabasz score. SIL score: Silhouette Coefficient score.

We focused on four drugs (Sorafenib, NVP-TAE684, PLX4720, and Etoposide) that exhibited lower AUROC values as reported in the original study. These drugs had drug-sensitive labels of bulk data in the GDSC^30^ database and the cell-level drug-sensitive labels were obtained in different ways. For the drug PLX4720 and Etoposide-affected single cells, cells from drug-untreated cell lines were considered sensitive, while cells that survived after drug exposure were considered resistant^45^. For drug Sorafenib and NVP-TAE684 affected cells, the cells’ sensitive labels were determined by the value of senescence-related (EpiSen) program scores which were proved to have a relation with drug responses previously^46^.

We compared our results with the baseline SCAD model, which took all genes’ expression values as input. We found that the scFoundation embedding-based model achieved higher AUROC values for all four drugs (Fig. 4B). Particularly for drugs NVP-TAE684 and Sorafenib, the scFoundation embedding-based model exhibited an improvement of above 0.2 in AUROC. We then employed the Spearman correlation as a metric to assess the agreement between the predicted probability of drug sensitivity provided by the model and the EpiSen score. As depicted in Fig. 4C, drug sensitivity for NVP-TAE684 and Sorafenib was positively and negatively correlated with EpiSen scores, respectively. The scFoundation embedding-based model yielded a Spearman correlation of 0.56 and -0.55 for the two drugs, while the baseline model achieved a Spearman correlation of only 0.24 and -0.06. These findings indicated that the embedding obtained from scFoundation facilitated drug sensitivity prediction in the original single-cell dataset with poor performance, and the embedding-based SCAD model had the potential to capture the signal of drug sensitivity biomarkers.

These results further motivated us to investigate whether the embedding itself was more informative than gene expression without the necessity for the SCAD model to extract the signal. We conducted principal component analysis (PCA) on the single-cell dataset SSC47, corresponding to drugs NVP-TAE684 and Sorafenib. By visualizing cells with different EpiSen scores on the first two principal components (PC) plots (Fig. 4D), we observed that compared to gene expression data, the first two PCs of the embeddings displayed less linear correlation, indicating the provision of richer information. Additionally, we used drug sensitivity as the label and computed the clustering performance of the embeddings and gene expression for both bulk and single-cell data. Calinski-Harabasz (CH) and SIL scores were used as metrics. The results for both single-cell and bulk data (Fig. 4E and Fig. S3) demonstrated that the scFoundation embedding better-grouped cells or bulk cell lines with the same drug response, compared to the gene expression baseline.

Overall, these findings highlighted that the unified embedding obtained from scFoundation aligned bulk and single-cell data into a unified representation space, and this condensed representation effectively facilitated the transfer of pharmacogenomics information from bulk cell lines to single-cell data.

### scFoundation predicts more accurate perturbation responses

Cellular response to perturbation is crucial for biomedical applications and drug design, as it helps identify gene-gene interactions across different cell types and potential drug targets^47,48^. Perturb-seq^49,50^ was a recently developed method for screening single-cell gene expression responses to several perturbations. Using these perturbation data resources to train models for modeling cellular response to perturbations is a central goal of computational biology^51–53^.

We combined the scFoundation with an advanced model GEARS^51^ for the perturbation prediction task. In the original GEARS model, a gene co-expression graph was combined with perturbation information to predict the post-perturbation gene expression. Each node in the co-expression graph represented a gene, with initially randomized embeddings, and edges connected co-expressed genes. This graph was not cell-specific and was shared across all cells. In our setting, we obtained gene context embeddings for each cell from the scFoundation decoder and set these embeddings as the nodes in the graph (Method). This resulted in a cell-specific gene co-expression graph for predicting perturbations (Fig. 5A).

**Figure 5.**
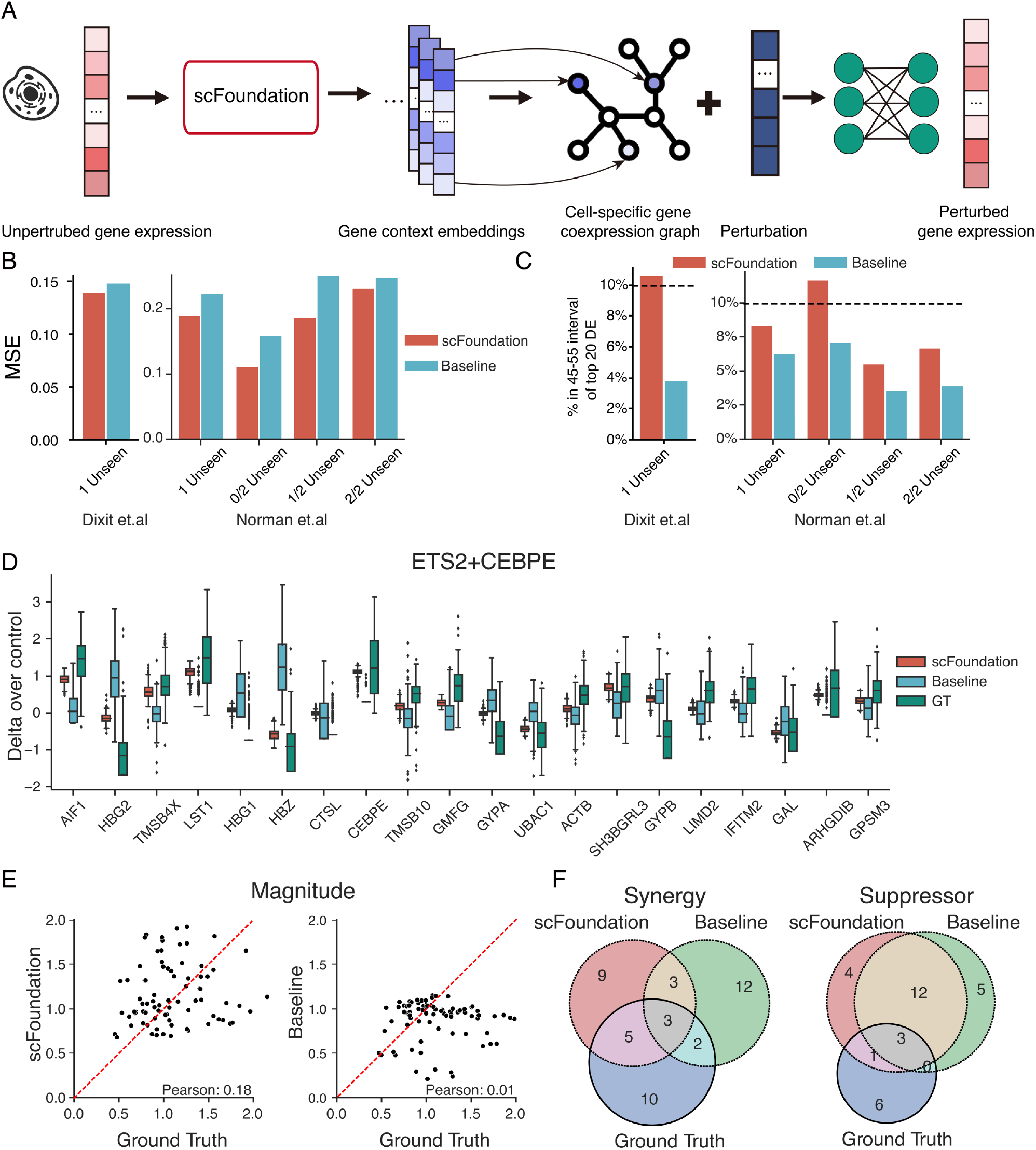
Perturbation prediction tasks using scFoundation gene context embeddings. A) Illustration of the perturbation prediction model based on cell-specific gene embeddings. B) Mean square error between predicted and ground truth post-gene expressions. Results given by the GEARS model using scFoundation cell-specific gene embeddings and gene expression are shown in red and blue, respectively. C) Proportion of predicted values within (+-5%) of the true mean expression value of the top 20 differentially expressed (DE) genes. The black dashed line represents the expected percentage (10%). Colors represent the same results as in panel A. D) Predicted gene expression over control for the top 20 most differentially expressed genes after a combinatorial perturbation (ETS2+CEBPE). The red and blue boxes indicate the gene prediction results by the GEARS model using scFoundation gene embeddings and pre-gene expression, respectively. The green box represents the ground truth post-gene distribution. E) Magnitude scores computed for all test perturbing combinations on the Norman dataset. Each dot represents a specific perturbing combination. The y-axis shows the magnitude score computed from the prediction results, while the x-axis represents the ground truth magnitude score computed using real post-gene expression. F) Top 20 perturbations with synergistic and suppressor gene interaction types identified using scFoundation and baseline methods. The Venn plot illustrates the relationship between the identified perturbation set and the verified perturbation set.

We trained and tested models on three perturbation datasets which were also used in the original study: the Adamson dataset^49^ with 87 1-gene perturbations, the Dixit^50^ dataset with 24 1-gene perturbations, and the Norman^54^ dataset with 131 2-gene perturbations and 105 1-gene perturbations. We reprocessed the gene numbers to the full gene length of 19,264, ensuring a comprehensive modeling of gene co-expression. For the 1-gene perturbation datasets, we leave a subset of perturbations for testing. For the 2-gene perturbation Norman dataset, we followed the original study to group 3 2-gene perturbations subsets for testing: 0/2, 1/2 and 2/2, where the first number indicated how many perturbations were not seen in the training set.

We computed the mean square error (MSE) of the top 20 differentially expressed (DE) genes between pre- and post-gene expression profiles to evaluate the models’ performance. The scFoundation-combined model achieved lower MSE values on all 1-gene perturbation datasets compared to the original GEARS baseline model (Fig. 5B and Fig. S4). For 2-gene perturbations, the model achieved the lowest MSE in the 0/2 unseen case and outperformed across all cases. We further examined the proportion of predicted values that fell within (+-5%) of the true mean expression value of the top 20 DE genes. The results in Fig. 5C demonstrated that the scFoundation-based model exhibited a higher percentage close to 10%, indicating that it provided a more reasonable distribution of post-gene expression values. We then showcased the 2-gene perturbation ETS2+CEBPE in Fig. 5D for an instance of prediction. The scFoundation-based model achieved a more accurate mean post-expression value and a distribution that closely aligned with the ground truth, outperforming the baseline model.

One application for predicting 2-gene perturbations was to classify 2-gene perturbation into different genetic interaction (GI) types. Here, we identified synergy and suppressor GI types by using the magnitude score (Methods). We first computed the PCC between predicted and ground truth magnitude scores of all test set 2-gene perturbations and found that the scFoundation-combined model achieved a higher PCC compared to the baseline (Fig. 5E). Next, we ranked the 2-gene perturbations according to the predicted magnitude score and regarded the top 20 and bottom 20 perturbations as potential synergy and suppressor GI types, respectively. The Venn plot in Fig. 5F revealed that the scFoundation-based model identified a higher number of true perturbations for both synergy and suppressor types.

These results highlighted the cell-specific gene context embeddings obtained from the scFoundation model served as valuable foundational representations for perturbation prediction. Furthermore, the analysis of 2-gene perturbations underscored the model’s capability to accurately classify different types of genetic interactions.

## Discussion

Recent breakthroughs in large-scale models demonstrated their ability to grasp complex natural language patterns through self-supervised pre-training. This motivated us to explore whether large-scale models can also be effective for learning the cellular and molecular “languages” of biology from single-cell transcriptomic data, which exhibit large data scales, complex biological patterns, diversity, and technical noises. We developed the xTrimogene architecture to enable non-destructive modeling of all protein-coding genes and continuous expression values in cells. Combining this architecture with our read depth-aware pre-training task, we presented scFoundation, the largest pre-training foundation model on the single-cell field, with 100 million parameters pre-trained on over 50 million single-cell data. We provided a comparison of the major features with the recently released similar models in Supplementary Table 1.

Our experiments showcased the model’s remarkable capabilities across various tasks. scFoundation can scale up effectively, surpassing other models in terms of parameter size. It can enhance read depth for gene expression without requiring additional fine-tuning, and can achieve superior gene expression restoration and cell clustering performance compared to other imputation methods. In the Cancer Drug Response task, the cellular embeddings derived from scFoundation significantly improved the prediction accuracy of drugs’ IC50 values. In the single-cell drug sensitivity task, scFoundation effectively integrated the unique characteristics of both bulk and single-cell data, achieving superior performance in drug prediction. In the perturbation prediction task, scFoundation can overcome the limitations of existing approaches by constructing cell-specific gene co-expression maps, yielding the most accurate perturbation predictions to date.

It is worth noting that scFoundation followed a similar training approach to the linear probing^55^ in all tasks, which added a task-specific predicting head after the pre-trained model output. The pre-trained model itself was not further fine-tuned, but the downstream task model was trained based on the obtained cell or gene context embeddings. This usage reduced computational and time costs for downstream users and offered flexibility in downstream model design. This paradigm allows scFoundation to better serve as a foundational model for a variety of downstream tasks in the field of single cell biology.

In the future, we will pre-train models with more parameters and larger datasets using our effective pre-training framework. Additionally, the availability of single-cell multi-omics data^56,57^ opens up new avenues for modeling cellular regulatory relationships and cell state transitions^58^. These data can be used to train multi-omics large-scale models and to provide deeper insights into cellular biological processes, towards modeling the multi-level complex laws of cells.

We expect that our pre-training architecture and the scFoundation model can serve as fundamental contributions supporting both large-scale biological models and a variety of downstream research. Our experiments as well as other recent reports suggest that such large biological models (LBM) pretrained on high-throughput single-cell data can bring revolutions for future biological and medical discoveries and applications.

## Methods

### Pre-training Data collection and preprocessing

#### Data collection

The human scRNA-seq public data were stored in the Gene Expression Omnibus (GEO) repository, human cell atlas, Single Cell Portal, EMBL-EBI, etc. There were also several studies to integrate human single cells from multiple resources, such as hECA^59^, DISCO^60^, etc. Each dataset in these databases was linked to a published study and thus had a corresponding DOI id. We manually collected scRNA-seq data from these databases and removed the dataset with duplicated id. Most of the datasets provide the raw count matrix. For the dataset with normalized expression profiles, we converted them back to the raw count form: We treated the smallest non-zero value in the original matrix as a raw count value of 1, all remaining non-zero values were divided by this smallest value and the integer part was taken. For the dataset with TPM or FKPM expression profiles that cannot be converted back to raw counts, we kept them unchanged.

We collected more than 50 million single cells across multiple organs (e.g. heart, kidney, brain) and tissues (i.e. connective tissues, epithelial tissue, muscle tissue, and nerve tissue). Our database also included cells from various diseases and various cancer dissection regions of different cancer types, aiming to cover all known possible gene expression profiles of the human single cells.

#### Gene symbol unification

We unified the gene symbols of all raw count gene expression matrices by using the gene symbol mapping reference provided by HUGO Gene Nomenclature Committee. We included human protein-coding genes and common mitochondrial genes, constituting a total of 19,264 genes. If some symbols were missing, we padded them with zero values.

#### Quality control

To filter extremely low-quality or damaged cells, we kept cells with over 200 genes expressed (i.e., expression vector with non-zero value count > 200) for pre-training by using the Seurat^61^ and Scanpy^62^ packages.

### scFoundation model architecture

We developed the xTrimoGene model as the backbone model of scFoundation. It had three modules: The embedding module converted scalar value into embeddings that were required for the transformer block; The encoder focused on the informative non-zero and non-masked expressed genes; And the decoder integrated information across all genes.

#### Embedding module

Given a cell’s gene expression value vector ***X***^*input*^ ∈ ℝ^*n*=19264^, the expression value 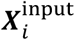 of gene *i* was a continuous scalar greater than or equal to zero. Unlike the previous language or recently developed single-cell transformer-based model, the embedding module directly converted the scalars into learnable value embeddings without any discretization. And then the value embeddings were added with gene embeddings to form the final input embeddings.

Specifically, the zero and non-zero values were converted in different ways. Zero value was directly converted into a randomly initialized embedding (with dimension *d*):

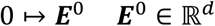

For the non-zero expression value, a look-up table ***T*** ∈ ℝ^*b*×*d*^ was randomly initialized, where *d* was the hidden embedding dimension and *b* was the pre-defined retrieved token number. We used *b* = 100 for our scFoundation model. Then for a non-zero expression value 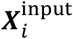 of gene *i*, it was first fed into a linear layer with leaky ReLU activation to get an intermediate embedding ***ν***_**1**_ :

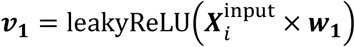

where ***w***_1_∈ ℝ^1×*b*^ was the parameter. The n another lin ear layer and a scaling factor *α* were used to further process ***ν***_**1**_ :

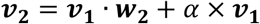

where ***w***_**2**_ ∈ ℝ^*b*×*b*^ and *α* were the learnable parameters. And the obtained vector ***ν***_**2**_ was normalized with the SoftMax function to get the attention score vector ***ν***_**3**_ ∈ ℝ^*b*^:

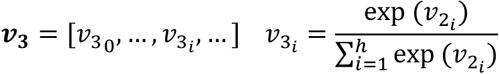

And the final value embedding ***E***_*i*_ ∈ ℝ^*d*^ of gene *i* was the weighted summation of all embeddings in the look-up table ***T***:

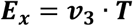

During pre-training, some expression value in the input gene expression ***X*** would be randomly replaced by the masked value *m*, and the masked value were corresponding to a mask embedding:

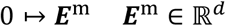

After converting scalar values into embeddings, an additional gene embedding was necessary to distinguish and indicate different genes. This design was similar to the positional embedding in the conventional language model^16^. We employed a gene look-up table ***T***^*G*^ ∈ ℝ^19266×*d*^ to retrieve embedding for all genes and two indicators T and S, where *d* is the same hidden dimension size as value embedding. For each gene *i*, the final input embedding 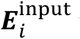 was the element-wise sum of the gene and value embeddings:

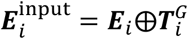

The input embeddings of all genes formed the final input tensor ***X***^*input*^ = [***E***_1_, ***E***_2_, …, ***E***_19 266_]^*T*^ ∈ ℝ^19266×*d*^.

#### Encoder

The encoder only processed the embeddings of non-zero and non-masked values (i.e. the expressed genes and two total count numbers) and so that the input length of the encoder was about 10% of the full gene length. Denote 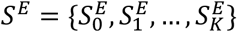 as the index set of non-zero and non-masked values with K elements, the input of encoder was defined as:

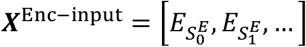

The design of encoder greatly reduced the required computational resources, making it possible for the encoder to employ a series of vanilla transformer blocks to capture gene dependency without any kernel or low-rank approximation. The outputs of encoder were intermediate embeddings ***X***^*inter*^:

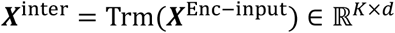

where Trm represents the transformer blocks. These intermediate embeddings had two usages: 1) they were then sent into the decoder with the zero and mask embeddings. 2) they could be pooled into cellular embeddings for downstream usages.

#### Decoder

To establish a transcriptome-wide gene regulation relationship, the zero-expressed genes should also be considered for recovering expression values at mask positions. The intermediate embeddings from encoder were concatenated with the zero and mask embeddings to form a decoder input tensor ***X***^Dec−input^ with full gene length:

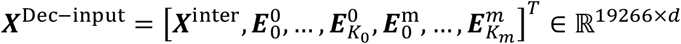

where *K*_0_ and *K*_*m*_ were the number of zero and masked embeddings, respectively. We used the kernel-based approximation transformer variant Performer^12^ as the transformer blocks in the decoder, since the attention calculation was challenging for long sequences^12,63^.

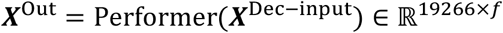

where ***X***^Out^ ∈ ℝ^19266×*d*^ was the output tensor of the decoder with dimension *f*. For predicting the expression value, the embeddings of T and S were dropped and a MLP was followed to project other *f* -dimension embedding to scalars. These scalars formed a prediction vector ***P*** :

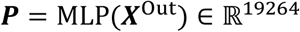

All parameters **Θ** = {***E***^0^, ***T***, ***w***_**1**_, ***w***_**2**_, *α*, ***E***^*m*^, ***T***, **Θ**_Encoder_, **Θ**_Decoder_, **Θ**_MLP_} were optimized during the pre-training. The detailed hyper-parameter setting of different models could be found in the Table S2.

### Read-depth-aware pre-training task

We trained our model with a read-depth-aware (RDA) gene expression prediction task. This task followed a similar self-supervised learning setting but with augmented input data. For each raw pre-training single-cell gene expression vector, we used a hierarchical Bayesian downsampling strategy to generate its low total count variant or unchanged profiles as the input vector. Then we defined two total count indicator T and S and set their values as the total count of the raw and input vectors, respectively. The raw and input vector were normalized and log-transformed.

Then we randomly masked the genes’ expressions of the input vector. In this study, we used 30% as our masking ratio for both zero and non-zero values. Then the partial masked input vector was concatenated with two total count indicator T and S and fed into the model. After getting the model-predicted raw gene expression, we conducted the regression loss on the masked genes between the predicted and the raw value. If the input vector was unchanged, the model learned to capture the relation between genes within a single cell. If the input vector was the low total count variant, the model learned the relationship between cells with different read depths.

#### Total count indicator

Besides gene expression values, there were two additional indicators S and T in our model input, indicating the value of the input and the output total count, respectively. At the pre-training stage, these two indicators were computed from the input and raw gene expression vectors. During the inference, the value of S was still computed from the sum of the input cell’s gene expression, but the value of T was set as the desired total count value (e.g., two folds of the input total count).

#### Hierarchical Bayesian downsampling strategy

For a single-cell gene expression vector ***X*** ∈ ℝ^19264^, we introduced a two-hierarchy Bayesian sampling to generate its corresponding input vector ***X***^*input*^. In the first hierarchy, a Bernoulli-distributed random variable *γ* was introduced to decide whether ***X*** would be downsampled. If *γ* equaled to 0, the input vector ***X***^*input*^ was the copy of ***X***; If *γ* equaled to 1, the input vector ***X***^*input*^ was the downsampled variant of vector ***X*** given by the second hierarchy. For those gene expression with total count lower than 1,000, we thought their quality were not good for downsampling and fixed *γ* = 0.

In the second hierarchy, we downsampled the raw count value of each gene via a binomial distribution:

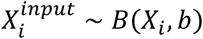

where 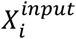 was the downsampled raw count value of gene i, *X*_*i*_ was the gene i observed raw count value and *b* was the downsampling rate. for each cell, all genes shared the same downsampling rate *b*. And thus the expectation of the downsampled cell’s total count satisfying:

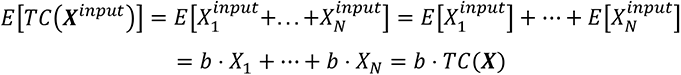

This design ensured that the expectation of the fold change between the raw and downsampled cell’s total count was fixed as 1*/b*. But for each gene in the ***X***^*input*^, the expression value is an observation of the random variable.

Further, we let the parameter *b* followed a beta distribution:

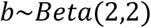

This prior distribution guaranteed that the different fold changes between input and output could be seen during training.

Overall, the input vector ***X***^*input*^ can be defined as:

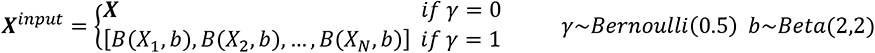

Then this input vector would be random masked and sent into the xTrimoGene model.

#### Mean square error (MSE) loss

Unlike other works that discretize expression values, our model operated on continuous raw gene expression values. So we used the regression loss as the loss function:

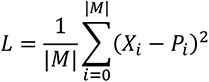

where M was a binary vector indicating which gene is masked (i.e. value 1 refers to be masked), *X*_*i*_ and *P*_*i*_ were the ground truth and predicted gene expression values, respectively. And | | was a 1-norm operator.

### Implementation

The attention mechanism in language modeling determined that increasing sequence length given near quadratic growth of time and space complexity. Even with the model architecture improvements which reduced the complexity from quadratic to linear, experiments with a large training corpus and large model parameter size still required a significant amount of resources. Therefore, time and resource-efficient training techniques were crucial for this work to collect abundant solid experimental results and supported training on 50M data.

HPC cluster infrastructure with NVIDIA Ampere GPU, NVLink for inter-GPU communication inside a server, and a high-speed interconnection network between servers, were utilized as a multi-node deep learning environment to optimize our experiment process.

Since half-precision operations could be executed on FP16 or BFLOAT16 Tensor Core which had 2 times more arithmetic throughput than TF32 on NVIDIA Ampere GPU, besides mixed-precision training decreased both the required amount of memory and the memory bandwidth consumption while maintaining model accuracy, our experiments were conducted with mixed-precision training strategy to gain time saving within a given amount of computational resource.

Distributed Data Parallelism was another widely used training strategy in our work to handle large corpus on HPC cluster, as our model architecture has greatly cut off the necessity of model parallelism in space and time complexity for long sequence modeling. Single Ampere GPU provided a sufficient amount of memory for one model replica of billions of parameters to perform forward and backward passes and gradient accumulation was used to increase the effective batch size to enhance large model training.

To achieve a larger model size without introducing model parallelism, ZeRO-DP stage two^64^ and checkpointing technique^65^ were experimentally verified reducing model state memory and residual state memory in our environment settings without expanding training time too much while introducing extra communication and computational costs.

For efficient and stable training of the model, we put the layer normalization inside the transformer block and thus the gradients were well-behaved at initialization^66^, reducing the hyperparameter tuning cost for the learning rate warm-up stage.

### Read depth enhancement analysis

For the gene expression prediction evaluation, we sampled 10,000 cells with a high total count (higher than 1,000) from 50 million single-cell data as the validation dataset. These 10,000 cells were excluded at the training stage. Then we used a binomial distribution to generate the low total count gene expression vector and fed it into our model. We only evaluate non-zero gene expression values considering that 0 expression values do not change in value after downsampling. In addition to using MSE as the evaluation metric, we also used the mean relative error (MRE), which can reflect the relative error:

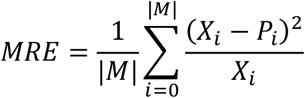

For the clustering analysis, we got the cell embeddings from scFoundation and scVI encoder. For others, we got the imputed gene expression profiles. All methods were used with the default parameter setting. Then we followed the SCANPY pbmc3k tutorial and got the cell cluster by the function ‘sc.tl.leiden’.

For the evaluation of the clustering results, we first used ARI and NMI as indicators to evaluate the degree of consistency between the clustering results obtained by different methods and the actual cell type labels. Considering that the acquisition of cluster labels will also be affected by the choice of the clustering algorithm, we used SIL as another evaluation indicator. Compared with ARI and NMI, SIL measures the aggregation degree of true cell type labels on the cell neighborhood maps given by different methods, and thus is independent of the choice of clustering algorithm, reflecting the intrinsic properties of cell representation.

### Downstream methods

We dumped the cell embeddings for DeepCDR and SCAD tasks, and gene embeddings for GEARS task. And we trained the downstream model based on these embeddings. All baseline models were trained with default parameters.

#### DeepCDR

We used the cell line and drug-paired data pre-processed by DeepCDR. The cell line data contains 697 gene expression profiles, and we aligned these genes with our unified list. The drugs were represented as graphs with consistent feature matrices and adjacent matrices sizes.

In total 223 drugs and 561 cell lines data from 31 cancer types were considered and 89,585 and 4,729 cell line-drug samples were used for training and test, respectively. For each cell line, we set both indicators S and T equal to the sum of all gene expression values. And we fed the non-zero gene expression values and two indicators into the model encoder and got the context embedding for each gene. The cell line embedding was obtained by the max-pooling operation for each embedding dimension across all genes.

We trained the baseline model by setting parameters “-use_gexp” as True and “-use_mut” and “-use_methy” as False. Then for the embedding-based model, we directly replaced the gene expression with the cell embedding and trained the DeepCDR with the same setting. Pearson’s correlation coefficient (PCC) was used as the evaluation matrix. For each gene, we computed the PCC between predicted IC50 and truth IC50 across all cell lines. For each cell line, we computed the PCC across all drugs conducted on this cell line.

#### SCAD

For training the baseline model, we used the processed data provided in their repository. Gene expression values were transformed into the z-score in the processed data. We used all genes and conducted the weighted sampling in the model training process, following the same experimental setting as the original SCAD study.

For training the embedding-based model, we used the normalized gene expression data. For bulk data, we set both S and T equal to the sum of all gene expression values. The same values of S and T would guide the model to keep the original cell line features. For single-cell data, we set the marker S to the sum of all gene expression values, and uniformly set the marker T to 10,000, which empirically is the maximum sequencing depth of a single cell. This setting made the model output consistent in read depth and was as close to Bulk data’s read depth as possible. Then the non-zero values of each bulk-level/single-cell gene expression sample and two indicators were fed into the encoder of the pre-trained model. The outputs were the context embeddings of genes for each sample. We found that the best performance can be achieved by concatenating the embeddings obtained in four ways. 1) the max-pooling operation for each embedding dimension across all genes. 2) the mean-pooling operation for each embedding dimension across all genes. 3) the context embedding corresponding to the indictor S. 4) the context embedding corresponding to the indictor T. These four types of embeddings built the new cell embeddings with 3072 dimensions. We used these cells’ embeddings to train a new SCAD model.

#### GEARS

We downloaded the raw gene expression data and unified the gene list to 19,264. We regenerated the gene co-expression network on each dataset. Then we trained the baseline model by setting epoch to 15 and batch size to 30. For the embedding-based model, we first set each cell’s T and S values equal to the its total count. Then the gene expression and these two indicators were fed into the model. We dropped the last MLP layer in our model and got the gene context embeddings from decoder. The cell-specific gene context embeddings were set as the node features of the co-expression graph. Then we trained the GEARS model’s parameter and froze others. We used the gradient accumulation technique to guarantee the same batch size as the baseline during training.

We followed the definition and metrics used in GEARS and Norman. We focused on the synergy and suppression gene intersection types since they were the most basic types. Identification of these two types was based on the magnitude score which measured the similarity between the two-gene perturbation and combining two single-gene perturbations. Specifically, let the mean change between post- and pre-A perturbed cells as *δg*_*a*_. A linear model was used to fit the effect of *δg*_*a*_, *δg*_*b*_, and *δg*_*a*+*b*_:

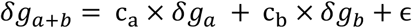

where ϵ captures the error in the model fit. We used the robust regression with a Theil-Sen estimator following the same procedure used in previous study^54^. Using the values of the coefficients, magnitude was defined as:

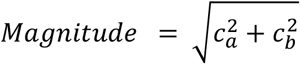

All test two-gene perturbations were ranked by magnitude score, with the top and bottom-ranked being considered synergistic and repressive types, respectively.

## Supporting information

Evaluation Metrics & Supplementary Figures & Supplementary Tables

## Data availability

All data used in this study are publicly available and the usages are illustrated in the Methods. The pre-training datasets were mainly downloaded from GEO (https://www.ncbi.nlm.nih.gov/geo/), Single Cell Portal (https://singlecell.broadinstitute.org/single_cell), HCA (https://data.humancellatlas.org/), EMBL-EBI (https://www.ebi.ac.uk/). The datasets used for downstream tasks can be downloaded from the following link: Baron dataset(https://github.com/mohuangx/SAVER-paper); Zheng68K dataset (https://www.dropbox.com/sh/w3yg2nucnng5v1u/AAAM8Ym_KU9XF4z51RT81eNEa?dl=0 processed by Wenpin et al.); Cancer drug response dataset (https://github.com/kimmo1019/DeepCDR); Single cell drug response classification dataset(https://github.com/CompBioT/SCAD); Perturbation dataset (https://github.com/snap-stanford/GEARS).

## Code availability

The code for producing the baseline results is in the downstream methods’ GitHub repository. The code for the scFoundation embedding-based model is publicly available at https://github.com/biomap-research/scFoundation.

## Acknowledgements

We thank Qijin Yin, Linlin Chao, Zhaoren He from Biomap, Yixin Chen, Chen Li, Haiyang Bian, Jiaqi Li, Tianxing Ma and Dr. Rui Jiang from Bioinfo Division, Tsinghua University for discussions and comments. This work was partially supported by National Natural Science Foundation of China (NSFC) (grants 62250005, 61721003), National Key R&D Program of China (grant 2021YFF1200901) and Tsinghua-Fuzhou Institute for Data Technology (TFIDT2021005).

## Author contributions

M.H., J.M., L.S. and X.Z. (Xuegong Zhang) conceived the study. M.H. Xin Z. (Xin Zeng) and Y.G. collected the downstream datasets involved in this article. Y.G. and L.S. developed data collection criteria and strategies for pre-training. M.H., J.G., Xin Z., C.L., T.W. and X.C. proposed the pre-training framework. M.H., J.G., Xin Z. and C.L. implemented and pre-trained the models. M.H. and J.G. benchmarked all methods. J.G., X.Z., C.L., T.W. X.C., J.M., L.S. and X.Z. provided a lot of advice on pre-training framework design and downstream tasks. M.H., J.G., J.M., L.S. and X.Z. wrote the manuscript. All authors read and approved the final manuscript.

## Additional information

### Supplementary information

Evaluation Metrics & Supplementary Figures & Supplementary Tables

