## Supplementary material for "Large Scale Foundation Model on Single-cell Transcriptomics": Evaluation Metrics & Supplementary Figures & Supplementary Tables

#### *Adjust Rand index*

Let C is a ground truth cell type assignment and K is the clustering, and *a* is defined as the number of pairs of elements that are in the same sets in C and in the same set in K, *b* is the number of pairs of elements that are in different sets in C and in different sets in K. The Rand index is given by:

$$\mathrm{RI} = \frac{a+b}{C_2^{n_{samples}}}$$

And the ARI is given by:

$$\mathrm{ARI} = \frac{\mathrm{RI} - E[\mathrm{RI}]}{\max(\mathrm{RI}) - E[\mathrm{RI}]}$$

where $E[\mathrm{RI}]$ is the expected Rand Index (RI) of random labelings.

#### *Normalized Mutual Information*

Assume two label assignments of the same $N$ objects, $U$ and $V$. Their entropy is the amount of uncertainty for a partition set is defined by:

$$H(U) = -\sum_{i=1}^{|U|} P(i)\log\big(P(i)\big) \quad H(V) = -\sum_{i=1}^{|V|} P'(i)\log\big(P'(i)\big)$$

where $P(i) = |U_i|/N$ is the probability that an object picked at random from $U$ falls into class $U_i$. Likewise for $P'(i) = |V_i|/N$. The mutual information $\mathrm{MI}(U,V)$ is defined as:

$$\mathrm{MI}(U,V) = \sum_{i=1}^{|U|}\sum_{j=1}^{|V|} P(i,j)\log\left(\frac{P(i,j)}{P(i)P'(j)}\right)$$

And the normalized mutual information is defined as:

$$\mathrm{NMI}(U,V) = \frac{\mathrm{MI}(U,V)}{\mathrm{mean}\big(H(U),H(V)\big)}$$

***Silhouette Coefficient score***

Let $a$ is the mean distance between a sample and all other points in the same class, and $b$ is the mean distance between a sample and all other points in the next nearest cluster. The Silhouette Coefficient $s$ for a single sample is then given as:

$$s = \frac{b - a}{max(a,b)}$$

And the Silhouette Coefficient score for a set of samples is given as the mean of each sample's Silhouette Coefficient score.

***Calinski-Harabasz score***

For a dataset of size $n$ which has $k$ cell types, the Calinski-Harabas score $s$ is defined as the ratio of the between-clusters dispersion mean and the within-cluster dispersion:

$$s = \frac{\mathrm{tr}(B_k)}{\mathrm{tr}(W_k)} \times \frac{n-k}{k-1}$$

where $\mathrm{tr}(B_k)$ is trace of the between group dispersion matrix and $\mathrm{tr}(W_k)$ is the trace of the within-cluster dispersion matrix.

### Supplementary Figures

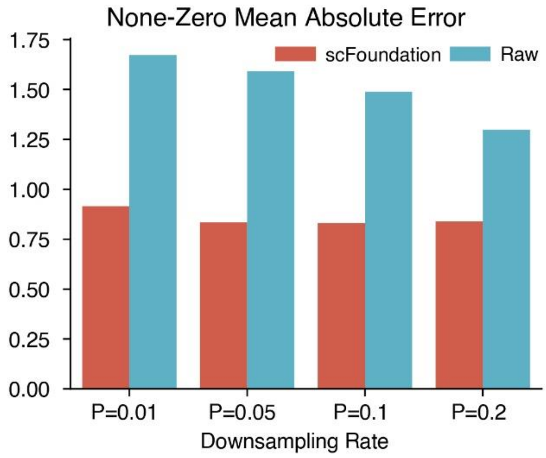


Figure S1. The mean absolute error among non-zero expressed gene.

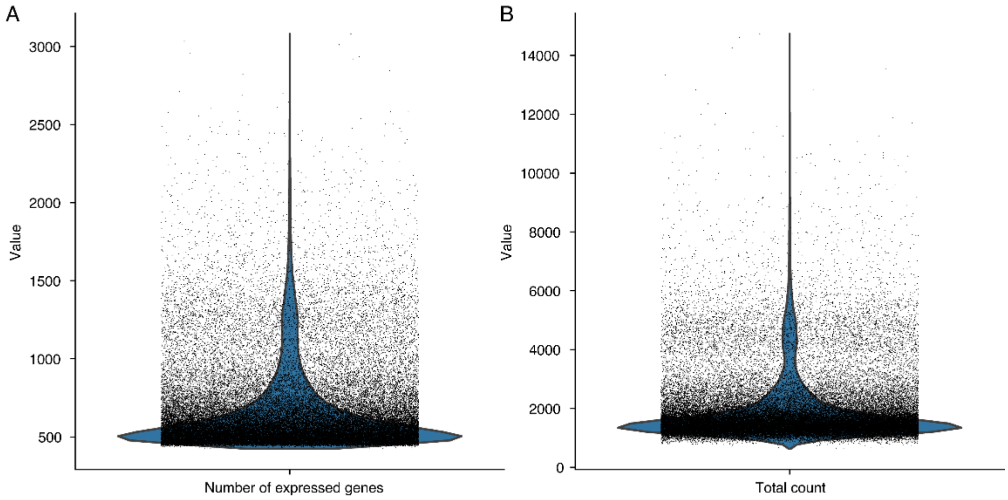


Figure S2. A) The distribution of expressed genes' number in a cell and B) the distribution of the total counts of cells on Zheng68K dataset.

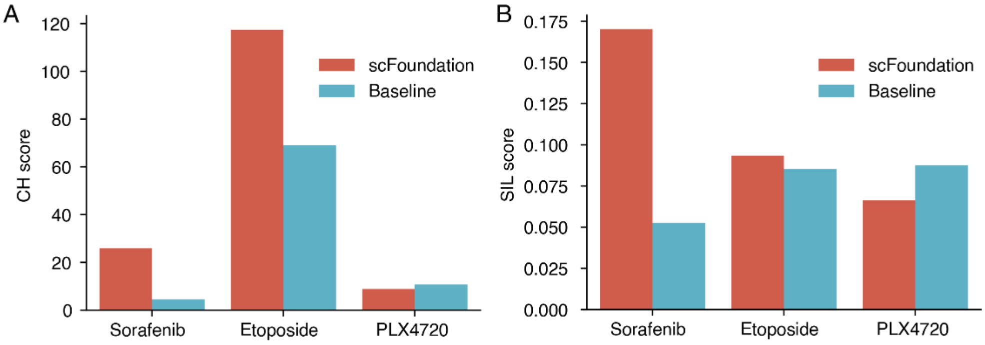


Figure S3. A) the Calinski-Harabasz score was computed on three drug-related single-cell datasets. B) the Silhouette Coefficient score computed on three drug-related single-cell datasets. Cells in each dataset are labeled by EpiSen-low and Epi-high groups.

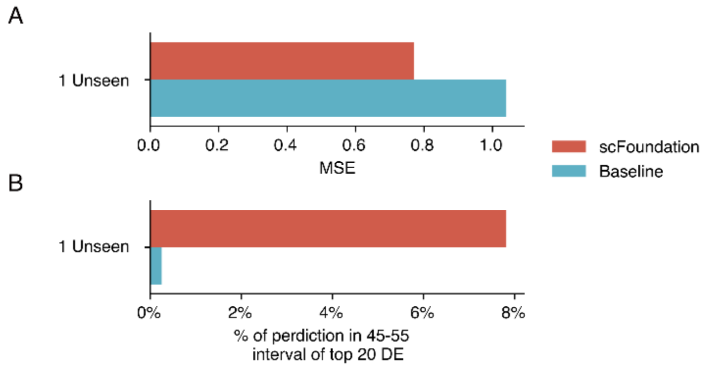


Figure S4. Performance evaluation on the Adamson dataset. A) Mean square error between predicted and ground truth post-gene expressions. B) Proportion of predicted values within (+-5%) of the true mean expression value of the top 20 differentially expressed (DE) genes. Results given by the GEARS model using cell-specific gene embeddings and gene expression are shown in red and blue, respectively.

### Supplementary Tables

Table S1: Model comparison

| Model | | **scFoundation** | scBERT | scGPT | Geneformer |
| --- | --- | --- | --- | --- | --- |
| Data | Size | **50M** | 1M | 10M | 30M |
| | Resources | Public data: HCA, GEO, hECA… | PanglaoDB | CELLxGENE portal | Public data |
| | Variability | **264 tissues** | 74 tissues | 6 tissues | 39 organs (tissue coverage not reported) |
| Pre-training | Model size | **~100M** | ~10M | ~50M | ~30M |
| | Model architecture | **Asymmetric Encoder-Decoder** | Encoder Only | Encoder Only | Encoder Only |
| | Design (Layer-Head-Dim) | Encoder Transformer: 12-12-768<br>Decoder Performer: 6-8-512 | Performer: 6-10-200 | Transformer: 12-8-512 | Transformer: 6-4-512 |
| | Input value | **continuous** normalized expression values | binned normalized expression values | binned normalized expression values | ranked normalized expression values |
| | Number of input genes (L) | **19,264** protein-coding or mt genes* | 16,906 genes | ~2,000 non-zero genes | 2,048 genes with different ranks |
| | token number (in 1 epoch training) | 50,000,000×19,264<br>= 963,200,000,000<br>**(~963B)** | 1,000,000×16,906<br>=16,906,000,000<br>(~16B) | 10,000,000×2,000<br>=20,000,000,000<br>(~20B) | 30,000,000×2,048<br>=61,440,000,000<br>(~61B) |
| | Model scalability | **Yes** | No (architecture limited) | No (length limited) | No (length limited) |
| | Training acceleration | MMF | vanilla Pytorch | | Huggingface (with Deepspeed) |
| | Pretraining Task | Read-depth aware masked value prediction** | Masked value prediction | Masked value prediction | Masked value prediction |

* Genes with zero expression values are also used in the training as their absence also provide information. See text for details.

** Using the gene expression of low read-depth cells to predict the expression of the high read-depth cells. See text for details.

Table S2: Size and hyper-parameters of different pre-trained models for testing scalability.

| Model Name | Parameter(M) | Encoder | | | Decoder | | |
|---|---|---|---|---|---|---|---|
| | | Depth | Heads | Dim | Depth | Heads | Dim |
| scFoundation (3M) | 3 | 4 | 2 | 128 | 2 | 2 | 128 |
| scFoundation (10M) | 10 | 4 | 8 | 256 | 2 | 4 | 256 |
| scFoundation (100M) | 100 | 12 | 12 | 768 | 6 | 8 | 512 |
